# RL-SKAT: An exact and efficient score test for heritability and set tests

**DOI:** 10.1101/140889

**Authors:** Regev Schweiger, Omer Weissbrod, Elior Rahmani, Martina Müller-Nurasyid, Sonja Kunze, Christian Gieger, Melanie Waldenberger, Saharon Rosset, Eran Halperin

## Abstract

Testing for the existence of variance components in linear mixed models is a fundamental task in many applicative fields. In statistical genetics, the score test has recently become instrumental in the task of testing an association between a set of genetic markers and a phenotype. With few markers, this amounts to set-based variance component tests, which attempt to increase power in association studies by aggregating weak individual effects. When the entire genome is considered, it allows testing for the heritability of a phenotype, defined as the proportion of phenotypic variance explained by genetics. In the popular score-based Sequence Kernel Association Test (SKAT) method, the assumed distribution of the score test statistic is uncalibrated in small samples, with a correction being computationally expensive. This may cause severe inflation or deflation of p-values, even when the null hypothesis is true. Here, we characterize the conditions under which this discrepancy holds, and show it may occur also in large real datasets, such as a dataset from the Wellcome Trust Case Control Consortium 2 (*n*=13,950) study, and in particular when the individuals in the sample are unrelated. In these cases the SKAT approximation tends to be highly over-conservative and therefore underpowered. To address this limitation, we suggest an efficient method to calculate exact p-values for the score test in the case of a single variance component and a continuous response vector, which can speed up the analysis by orders of magnitude. Our results enable fast and accurate application of the score test in heritability and in set-based association tests. Our method is available in http://github.com/cozygene/RL-SKAT.

## 1 Introduction

The variance component model is a well established statistical framework used in many scientific fields. Testing for an association between several explanatory variables and a univariate response produces a variety of useful applications. For example, in metagenomics, an association is tested between a phenotype (e.g, body mass index, blood glucose levels, blood lipid levels, etc.) and the relative abundance counts of the measured species [1].

In statistical genetics, testing for an association between a set of genetic markers and a phenotype, such as a disease or a trait, is a fundamental task. Since studies to detect genetic signals are often underpowered, even with large datasets becoming available, the common approach to help alleviate this issue is grouping together genetic markers and testing them jointly. Grouping genetic markers is commonly implemented under the framework of variance component models. In addition to association testing, this framework can be used to answer several questions, such as estimation of the underying heritability of a phenotype [2]; estimating the uncertainty of such estimation [3-5]; phenotype prediction [6], and more.

We consider two main scenarios in which such tests are performed: (i) a single phenotype, many sets of markers; (ii) many phenotypes, a single set of markers. Scenario (i) is common in set-testing, where relatively few markers are tested jointly. This is particularly useful in the case of rare variants, which are increasingly available for study using sequencing technologies, and which consistute a large part of human genetic variability. In such studies, a single phenotype is often tested against several sets of markers (for example, all rare variants in a single gene), because single-marker tests are often underpowered. Scenario (ii) occurs when studying heritability, defined as the proportion of phenotypic variance explained by genetics. Here, the tested markers are commonly the entire set of genotyped or sequenced single-nucleotide polymorphism (SNP) variants, or large portions of the genome (defined by, e.g., chromosome or functional annotation), and they are often tested against many (e.g., thousands) of phenotypes. Such phenotypes could be expression profiles of genes [7-9], methlyation levels across of various methylation sites in the DNA [10, 11] or neuroimaging measurements [12, 13].

Within the variance components framework, a common approach for association testing is the score test, which is based on calculating the derivative of the likelihood function at the point corresponding to zero association, and testing if it is significantly nonzero. Compared with another popular alternative, the generalized likelihood ratio (LR) test, the score test is often advantegeous as it requires parameter estimation only for the null model, whereas the LR test requires parameter estimation for both the null and the alternative model. Additionally, the score test is the locally most powerful test; see [14] for a discussion.

The Sequence Kernel Association Test (SKAT) [15] has become the standard score-based test in statistical genetics and in metagenomics [1], in large part due to its computational tractability. One of its merits is that it does not rely on the asymptotic distribution of the score test statistic, instead specifying a non-asymptotic distribution for the statistic under the null hypothesis of no association. However, it has been observed that this distribution may be inaccurate. In the SKAT-O extension [16], a resampling-based moment-matching correction is suggested. An adaptive permutation testing procedure is suggested in [17]. Chen et al. [18] provide a method for calculating exact p-values; however, their method may be significantly slower than that of SKAT, as it requires the eigendecomposition of a full rank square matrix, whose computational complexity is typically cubic in the sample size, for each distinct response variable (e.g., phenotype) or each set of explanatory variables (e.g., SNP set). Finally, in these works, it is reported that this discrepancy occurs mainly in studies having a small sample size, and it is currently unclear to which extent the p-values of SKAT are calibrated for large sample sizes.

Here, we undertake a thorough analysis of the null distribution of the score test statistic, and its discrepancy under the SKAT approximation. We suggest a practical way to quantify this discrepancy, and show that such discrepancies may occur even at large sample sizes. We show that a discrepancy is expected when the number of markers is comparable to or larger than the number of individuals, and when the individuals are relatively unrelated. In particular, in addition to such inaccuracies occuring in tests of sets of rare-variants in small samples, we conclude that they may also occur in large scale heritability studies. We further suggest a computational method, Recalibrated Lightweight SKAT (RL-SKAT), that allows exact p-value computation while maintaining computation time as in SKAT; in particular, for multiple phenotypes tested against the same marker set, only a single eigendecomposition is required. Finally, we demonstrate and validate our results on two real datasets, a large dataset from the Wellcome Trust Case Control Consortium 2 [19] (WTCCC2) study and the Cooperative health research in the Region of Augsburg (KORA) study [20] dataset.

## 2 The score test under the variance components model

We begin by reviewing the score test, as defined by the SKAT method [15] (see also the Supplemetary Information of [14] for an excellent review). We focus here on continuous phenotypes, and on the case of a single variance component; for other cases, see the discussion below.

### 2.1 The variance components model

We consider the following standard variance components model (see [21] for a detailed review). Let *n* be the number of observations and **y** be a *n* × 1 vector of responses. Let **X** be a *n* × *p* design matrix of *p* covariates, associated with fixed effects (possibly including an intercept vector **1**_*n*_ as a first column, as well as other covariates) and let ***β*** be a *p* × 1 vector of fixed effects. Finally, let **K** be a kernel matrix, which, in a kernel-based method such as SKAT, can be taken to be any symmetric positive-definite matrix that encodes similarity between individuals. Then, **y** is assumed to follow:

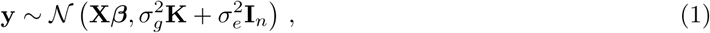

The fixed effects ***β*** and the coefficients 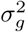 and the coefficients 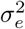 are the parameters of the model.

In the context of statistical genetics, **y** is a vector of phenotype measurements for each individual and **X** is a matrix of covariates (often including an intercept, sex, age, etc.). Let **Z** be a *n* × *m* standardized (i.e., columns have zero mean and unit variance) genotype matrix containing the *m* SNPs we test. The common choice for **K** is a weighted dot product of the genetic markers [22]; formally, define **K** = **ZWZ**^⊤^, where **W** is a non-negative *m* × *m* diagonal matrix assigning a weight per SNP. A standard choice is the uniform **W**_*i,i*_ = 1/*m* (see [15] for a discussion). The narrow-sense heritability due to genotyped common SNPs is defined as the proportion of total variance explained by genetic factors [23]:

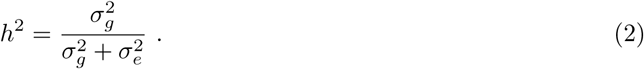

### 2.2 The score test

Under the above model, evaluating whether the tested covariates influence the response, while adjusting for additional covariates, corresponds to testing the null hypothesis 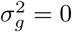. SKAT tests this hypothesis with a variance component score test in the corresponding mixed model. Specifically, the score statistic in the single-kernel case is obtained from the derivative of the restricted likelihood, discarding terms which are constant with respect to **y** [14]:

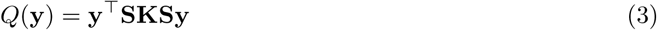

where **S** = **I**_*n*_ − **X**(**X**^⊤^**X**)^−1^**X**^⊤^ is the projection matrix to the subspace orthogonal to the covariates **X**. For clarity of presentation, we will divide the statistic by 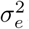. Then,

#### Proposition 1.

*Let* {*ϕ_i_*} *be the eigenvalues of* **SKS**^⊤^*and be* 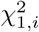 *are i.i.d. random variables distributed chi-square with one degree of freedom. Then*,

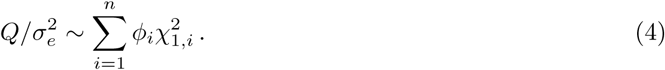

The proof of Proposition 1, as well as all proofs below, are deferred to Appendix A.

### 2.3 The exact distribution of the score test statistic

The above derivation is exact whenever 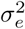 is known. However, in practice, 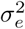 is not known and needs to be estimated from the data; most often, from the single response vector we are testing. In practice, 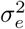 is replaced with its restricted maximum likelihood (REML) estimate. The REML estimate is simply the corrected mean of the squared entries of the phenotype, after regressing out the covariates and using **S**^⊤^**S** = **S**:

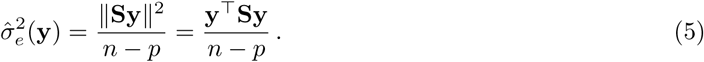

We note that sometimes the ML estimate **y**^⊤^**Sy**/*n* is used, or just **y**^⊤^**Sy**; as this only introduces a multiplicative constant, we use the unbiased REML estimate for simplicity of presentation later. The statistic *Q* and 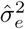, are in fact dependent random variables. Therefore, the assumed distribution of 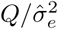 (described in Proposition (1)) does not hold when substituting 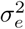 with its estimate, 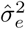. In [15, 24-26], this subtitution is justified by the claim that the (restricted) ML estimator 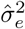 is consistent, and may therefore be substituted by its true value for a sample size *n* large enough. However, this argument does not take into consideration the dependency between *Q* and 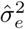. Also, as shown below, this distribution might not hold in realistic settings. In Chen et al. [18], this discrepancy is reported for small samples, and an exact distribution is derived for the statistic 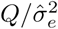, and for any *n*, **K** and **X**, which we review here:

#### Proposition 2.

*The distribution of* 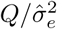 *may be modeled as a ratio of quadratic forms of normal variables. In particular, if* 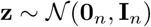*, then*

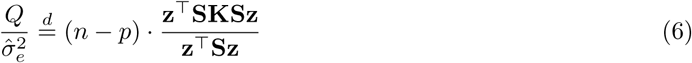

## 3 Assessing the discrepancy

While noted in the literature [1, 18], the above discrepancy is reported for small samples only. However, as we show now, it may occur also when the number of individuals is large. We give a qualitative measure for when to expect large discrepancies between the asymptotic approximation of a weighted mixture of chi-squares and the exact distribution.

In the Appendix, it is shown that the distributions of 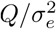 and 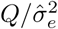 have the same means, but that 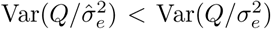, i.e. the latter having a smaller variance. We can further quantify the ratio between the variances as an indicator to the discrepancy between the distributions.

### Proposition 3.

*Denote the eigenvalues of* **SKS** *by ϕ*_1_*, …, ϕ_n_ and note that there are at most n* − *p non-zero eigenvalues ϕ_i_. Denote the first two sample moments of the eigenvalues by* 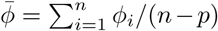 *and* 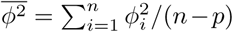*. Denote the empirical variance of the eigen values by* 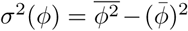*. Then*,

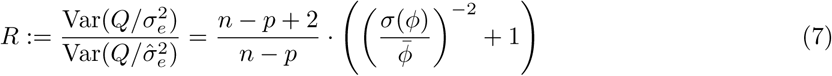

The expression 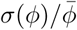 is the (sample) coefficient of variation (CV) of the eigenvalues − a unitless, relative measure of their dispersion. Therefore, the ratio becomes larger when the CV is smaller. Also, as noted above, since the approximation wrongly ignores the dependency between the statistic *Q* and 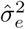, we expect the discrepancy to grow larger as the correlation between *Q* and 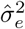 increases. We therefore examine this correlation as an additional measure of this discrepancy.

### Proposition 4.

*Let* 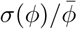 *be the coefficient of variation (CV) of the eigenvalues as above. Then*,

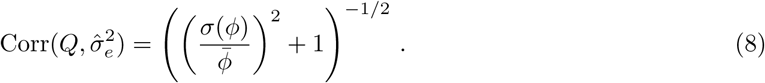

This again demonstrates that CV affects discrepancy – the correlation becomes stronger when the CV is smaller. When *CV* ≪ 1, for example when **K** ≈ **I**_*n*_, we have *R* ≫ 1 and 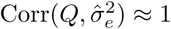. Conversely, when *CV* ≫ 1, we have *R* ≈ 1 and 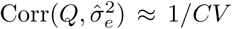. This also gives the variance ratio as the function of the correlation as

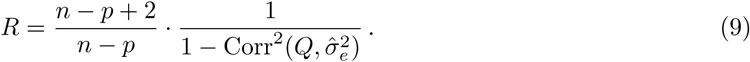

To summarize, the discrepancy is strong when the eigenvalues are more uniformly dispersed, and is weak when they have large variability.

**Examples.** We now employ Propositions 3 and 4 to analyze several interesting examples in a genetic context. For simplicity, in the following, we use **X** = **0**, so that *p* = 0 and **S** = **I**_*n*_.

- **Completely unrelated cohort.** Suppose the cohort contains completely unrelated individuals; then, **K** = **I**_*n*_. Thus, *ϕ*_1_ = *…* = *ϕ_n_* = 1, so *R* = ∞, 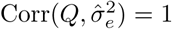, and 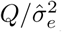 is the constant *n*. Compare this to the case where 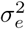 is known; then, it can be easily seen that 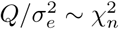. Therefore, the mean is the same but the variance vanishes completely.
- **Rank-one kinship matrix.** Consider the case of a simple burden test [16]: If we assume the random effects **s** of all SNPs are identical, the burden test becomes equivalent to the score test with **K** = **uu**^⊤^, where **u** = **Z1**_*m*_. Alternatively, consider the extreme case, where all the individuals are identical - **K** = **11**^⊤^ (while unlikely in human, this could be approximately true in studies of plants, yeast, etc.). In both these cases, there is a single nonzero eigenvalue: *ϕ*_2_ = *…* = *ϕ_n_* = 0, which gives *R* ≈ 1 and 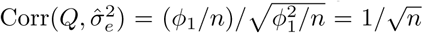; that is, with large enough sample size, we expect the correlation to be effectively zero, and the SKAT mixture approximation to hold well.
- **A full rank kinship matrix.** Assume the matrix **Z** contains *m > n* SNPs in linkage equilibrium, where each column was mean-centered and normalized to have unit variance. Choosing the linear kernel **K** = **ZZ**^⊤^ */m*, we follow [27] in modeling **Z** as a matrix of random standard normal variables, from which it follows that **K** is a Wishart matrix. The limit distribution of the density of the eigenvalues of **K** is specified by the Marčhenko-Pastur distribution [28], with its first two moments known to be 1 and 1 + *n/m*. Under this approximation, 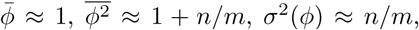, *R* ≈ (*n* − *p* + 2)/(*n* − *p*) · (1 + *n/m*)/(*n/m*) and 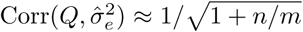. When *m* ≫ *n*, as is often the case, *R* ≫ 1 and 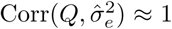. This shows that for a large class of kinship matrices, we would expect the SKAT mixture approximation to hold poorly.
- **A SNP set.** Now, consider the case of set-testing, where **Z** is a normalized matrix of *m < n* SNPs in linkage equilibrium. Following the modeling above, we have again *R* ≈ (*n* − *p* + 2)/*n* − *p*) · (1 + *n/m*)/(*n/m*) and 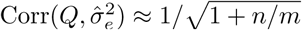; when *m* ≪ *n*, *R* ≈ 1 and 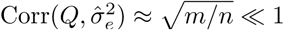, and thus expecting a good approximation by the mixture. This perhaps shows why the SKAT mixture approximation was considered good in the context of set-tests, when few variants or a large sample is considered. This also shows why, in small samples, the mixture is expected to be a poor approximation.

## 4 Calculating p-values

We now describe how to efficiently calculate p-values for the distribution of the statistic 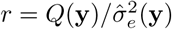 calculated from the data; that is, given an observed statistic *r*, what is 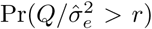 under the null? We review the result in Chen et al. [18]:

### Proposition 5.

*Let r be the observed value of the statistic. Denote by* 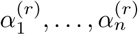 *the eigenvalues of* **SKS** − *r*/(*n* − *p*) · **S***. Then*,

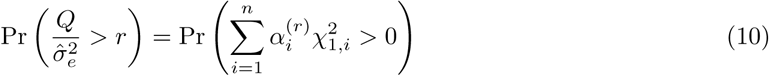

*where* 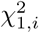 *are i.i.d. random variables distributed chi-square with one degree of freedom.*

However, this condition requires us to calculate the eigenvalues of **SKS**−*r*/(*n*−*p*)·**S** for each new value *r*, which, naively, has a complexity of *O*(*n*^3^). We consider two scenarios where this is problematic. First, in many heritability studies, we wish to test the heritability of many (e.g., thousands) of phenotypes, all relative to the same kernel or kinship matrix (see above). For each phenotype **y**_1_*, …*, **y**_*N*_, we calculate its score test statistic *r_i_*. For p-value calculation, we need to compute the eigendecomposition of **SKS** − *r_i_*/(*n* − *p*) · **S** for each observed statistic *r_i_*, which is a significant computational burden.

A second problematic scenario is of an association study of a single phenotype with many sets of SNPs, e.g. rare variants. Choosing a weighted linear kernel as in SKAT [15], we have 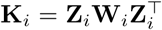 for each set. As **K**_*i*_ changes with each test, in principle, we need to perform a costly *O*(*n*^3^) eigendecomposition for each matrix **K**_*i*_. However, a significant computational saving is gained due to the fact that the nonzero eigenvalues of 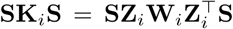 are the same as those of 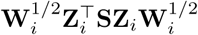, which is an *m* × *m* matrix [14]. As the number of tested SNPs *m* is often small, calculating the eigenvalues of this matrix instead is significantly faster, taking only *O*(*m*^3^), with matrix construction taking only *O*(*n*(*m* + *p*)^2^) (see [14]). However, with the exact approach, we need to calculate the eigenvalues of **SK**_*i*_**S** − *r_i_*/(*n* − *p*) · **S** instead of **SK**_*i*_**S**. Even when **K**_*i*_ is low rank, the matrix **SK**_*i*_**S** − *r_i_*/(*n* − *p*) · **S** may be close to full rank, so another approach is needed.

The following characterizes the eigenvalues of **SKS** − *r*/(*n* − *p*) · **S** given the eigenvalues of **SKS**:

### Proposition 6.

*Let r be the observed score test statistic. Denote by ϕ*_1_*, …, ϕ_n_ the eigenvalues of* **SKS***. Denote the column space of a matrix* **A** *by* col(**A**)*, its null space by* ker(**A**)*. Then*,

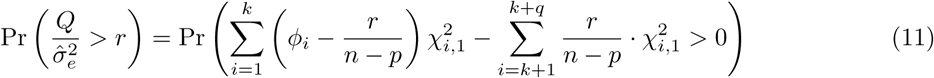

*where k* = rank(**SKS**) *is the number of nonzero eigenvalues ϕ_i_, q* = dim(ker(**SKS**) ∩ col(**S**))*, and* 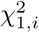 *are i.i.d. random variables distributed chi-square with one degree of freedom, i* = 1*, …, k* + *q.*

Proposition 6 shows that calculating the p-value amounts to evaluating the cumulative distribution function (cdf) of a certain weighted mixture of chi-square distribution at 0. This can be done rapidly using the Davies method [29], which is based on the numerical inversion of the characteristic function and runs in *O*(*n*) complexity, or using other methods [30].

It remains to calculate *k* and *q*. Naively, this can be done in *O*(*n*^3^), and when the same kernel is used with many phenotypes, it is a single preprocessing step. However, when the number of SNPs used to construct the kernel and the number of covariates are small, these quantities can be calculated much faster:

### Proposition 7.

*Suppose* **K** = **ZWZ**^⊤^*, and let k* = rank(**SKS**) *and q* = dim(ker(**SKS**) ∩ col(**S**))*. Then, k and q can be calculated in complexity O*(*n*(*m* + *p*)^2^).

Most commonly, *k* = min(*m, n*) − 1. When the number of SNPs *m* and the number of covariates *p* are small, the computational saving is substantial.

### 4.1 Performance summary

We summarize the results above in Table 1. We compare our method, RL-SKAT, with the SKAT formulation and the correction of [18] using the naive implementation of Proposition 5, as implemented by the MiRKAT software package [1]. The two scenarios discussed are those of a heritability study (same **K** with many responses **y**_*i*_) and SNP set-testing (many low rank **K**_*i*_). In all methods, a preprocessing step of calculating **X**^+^ and {*ϕ_i_*} is required. In a heritability study, calculating the statistic 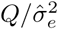 amounts to evaluating two quadratic forms in *O*(*n*^2^). Compared to RL-SKAT, MiRKAT requires a full *O*(*n*^3^) eigendecomposition for each **y**_*i*_. For a set-testing study, these quadratic forms can be calculated in *O*(*n*(*m* + *p*)) due to the low rank of **K**_*i*_. Again, MiRKAT requires a full *O*(*n*^3^) eigendecomposition, compared to the *O*(*n*(*m* + *p*)^2^) procedure described in Proposition 7.

**Table 1:** Performance summary. Comparison of the different approaches for p-value calculation discussed. RL-SKAT achieves accuracy while remaining computationally efficient.

| Scenario | Algorithm | Exact? | Preprocessing | Calculating $Q/\hat{\sigma}_e^2$ | Calculating p-value |
| --- | --- | --- | --- | --- | --- |
| Heritability | SKAT | Approximate | $O(np^2 + n^2p + n^3)$ | $O(n^2)$ | $O(n)$ |
| | MiRKAT | Exact | $O(np^2 + n^2p)$ | $O(n^2)$ | $O(n^3)$ |
| | RL-SKAT | Exact | $O(np^2 + n^2p + n^3)$ | $O(n^2)$ | $O(n)$ |
| Set-testing | SKAT | Approximate | $O(np^2 + nmp + nm^2)$ | $O(n(m+p))$ | $O(n)$ |
| | MiRKAT | Exact | $O(np^2 + nmp)$ | $O(n(m+p))$ | $O(n^3)$ |
| | RL-SKAT | Exact | $O(np^2 + nmp + n(m+p)^2)$ | $O(n(m+p))$ | $O(n(m+p)^2)$ |

## 5 Real data study: WTCCC2 and KORA

We now demonstrate our results on two datasets: a dataset from the Wellcome Trust Case Control Consortium 2 [19] (WTCCC2) study and the Cooperative health research in the Region of Augsburg (KORA) study [20]. A full description of data preprocessing is given in Appendix B.

### 5.1 A simulation study using WTCCC2 data

We first analyze data with real genotypes from the WTCCC2 Multiple Sclerosis dataset, and simulated phenotypes. We used the same data processing described in [31], resulting in *m*=360,556 SNPs for *n*=13,950 individuals. We constructed the kinship matrix by a standard, uniformly weighted linear kernel. We sought to demonstrate the discrepancy between the true null distribution and the chi-square weighted mixture distribution. Following Proposition 4, we calculated the correlation to be 0.886 and variance ratio to be *R* = 4.69, indicating that a large discrepancy is possibly expected. To verify this, we simulated 10,000 random phenotypes, where each phenotype is a vector of i.i.d. standard normal variables. We tested whether the variance component is significantly greater than zero, and calculated their p-values under the assumption of either of the two distributions. In Figure 1, we show the quantile-quantile plots for the two sets of p-values. As evidenced, using the SKAT mixture distribution results in a severe deflation of small p-values, while using the correct distribution as in Proposition 1 results in an accurate p-value distribution. This shows that even for large sample sizes (*n*=13,950), such a discrepancy is possible.

**Figure 1:**
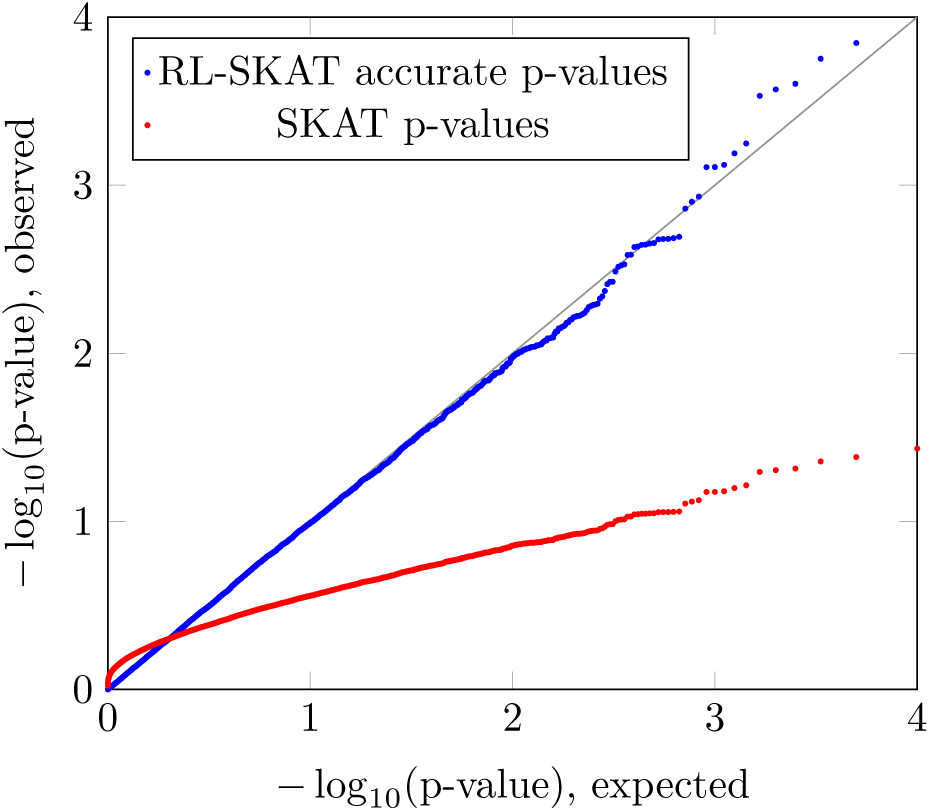
Statistic distribution. Results of the WTCCC2 data analysis, presented by quantile-quantile plots of the − log_10_(*p*)-values for heritability significance of 10,000 random phenotypes drawn under the null distribution. Significant deviation from the black line indicates a deflation arising from an inaccurate null distribution. Calculation under the assumption of a weighted mixture of chi-square distributions, gives deflated p-values and potentially creating false negatives. Using the correct distribution, as implemented in RL-SKAT, results in calibrated p-values.

### 5.2 Testing for heritable methylation sites in the KORA dataset

The longitudinal KORA study consists of whole-blood methylation levels and genotypes of *n*=1,799 individuals. The phenotype is the proportion of methylated samples at a specific site, averaged across DNA samples of an individual. The study consists of independent population-based subjects from the general population living in the region of Augsburg, southern Germany [20]. Whole-blood samples of the KORA F4 study were used as described elsewhere [32]. In summary, a total of 431,366 methylation site phenotypes, and 657,103 SNPs, were available for analysis. The correlation as in Proposition 4 is 0.976 and the variance ratio is *R* = 22.01, indicating again that a large discrepancy is expected. We performed a heritability study of multiple phenotypes with the same kinship matrix, by testing the heritability of the *N* =43,140 methylation sites on chromosome 1. As it is common for a methylation site to be correlated with its surrounding SNPs [33-35], we avoided such *cis* effects by using a kinship matrix constructed from the *m*=604,170 SNPs on all chromosomes other than 1. The kinship matrix is constructed by a standard, uniformly weighted linear kernel. For covariates, we used **X** consisting only of an intercept vector. Again, we calculated p-values under the assumption of the two distributions. We note that it has been shown that some methylation site profiles often display significant heritability, while others do not; thus, both significant and insignificant p-values are expected [36].

In Figure 2 we show the histograms of the log_10_ of the p-value of all the considered phenotypes. The two histograms are indeed very different; p-values calculated using the inaccurate SKAT mixture distribution indicate that the heritability of almost all sites is considered insignificant; for example, using a Bonferroni threshold of 0.05 · 1/43140 ≈ 10^−6^, only 8/43,140 sites are significant. In light of the results above, it is reasonable to suspect that p-values of many heritable phenotypes are deflated, thus causing false negatives. The p-values distribution has a peak around 0.5, likely an artifact of the inaccurate calculation method. In comparison, p-values calculated by RL-SKAT do not exhibit such a peak. They are significantly smaller, and using the same Bonferroni threshold, we now find 319/43,140 significant sites. Indeed, a simulated power study of both approaches under varying degrees of true underlying heritability validates that the inaccurate approach results in a severe decrease in power (Figure 3). We conclude that in this dataset, using the SKAT distribution for p-value calculation is highly problematic.

**Figure 2:**
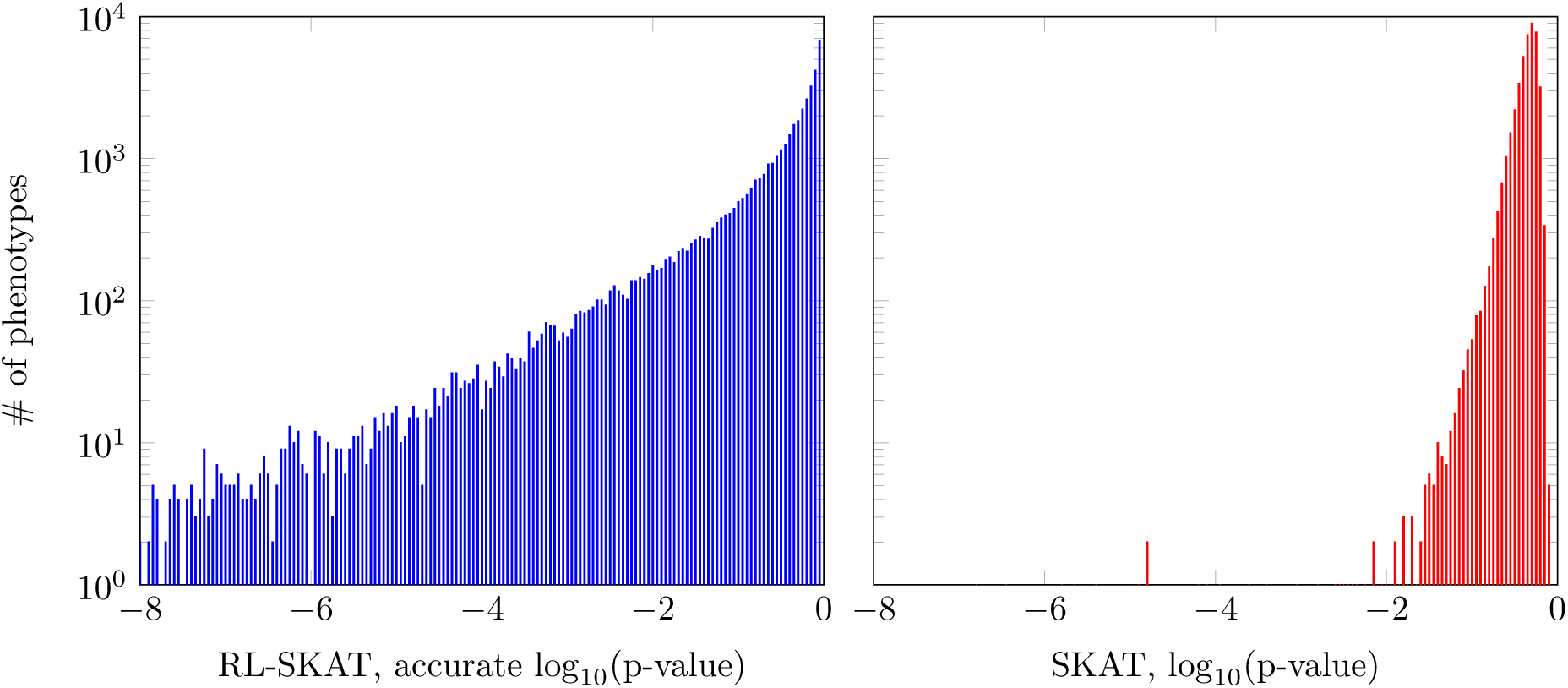
Heritability study. Histograms of the p-values of the studied phenotypes in the KORA dataset, as calculated by the accurate method (left) and the inaccurate method (right). Histograms are shown in log-scale, and are capped at *p* = 10^−8^ for clarity of presentation. SKAT tends to severely deflate p-values which are small according to the accurate calculation, leading to a severe loss of power.

**Figure 3:**
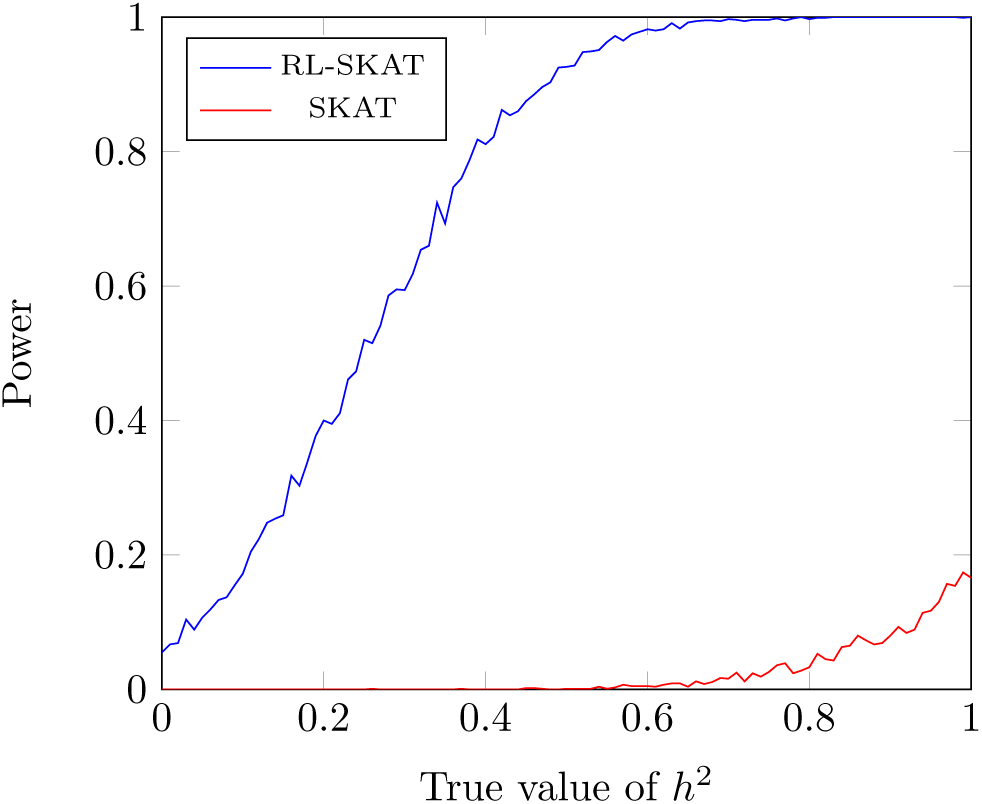
Power study. The power of the accurate approach and SKAT is shown for p-value threshold of *p* = 0.05, for the KORA dataset, on 10,000 simulated phenotypes with varying degrees of true underlying heritability. SKAT is seen to be severely underpowered.

### 5.3 Benchmarks

Finally, we benchmarked the methods discuss here on the KORA dataset under the two above scenarios, on a 64G, 2.2GHz Linux workstation, using our implementation in the Python language. We verified that the relevant part of our implementation is equivalent to MiRKAT and has a very similar running time. For the scenario of heritability testing, we calculated the p-values of 1000 phenotypes with the kinship matrix. For the scenario of set testing, we used 1000 sets of 100 SNPs each. The results are summarized in Table 2; as expected, the computational savings are very significant, achieving a speedup of more than two orders of magnitude. We expect the speedup to be even more significant for larger datasets.

**Table 2:**
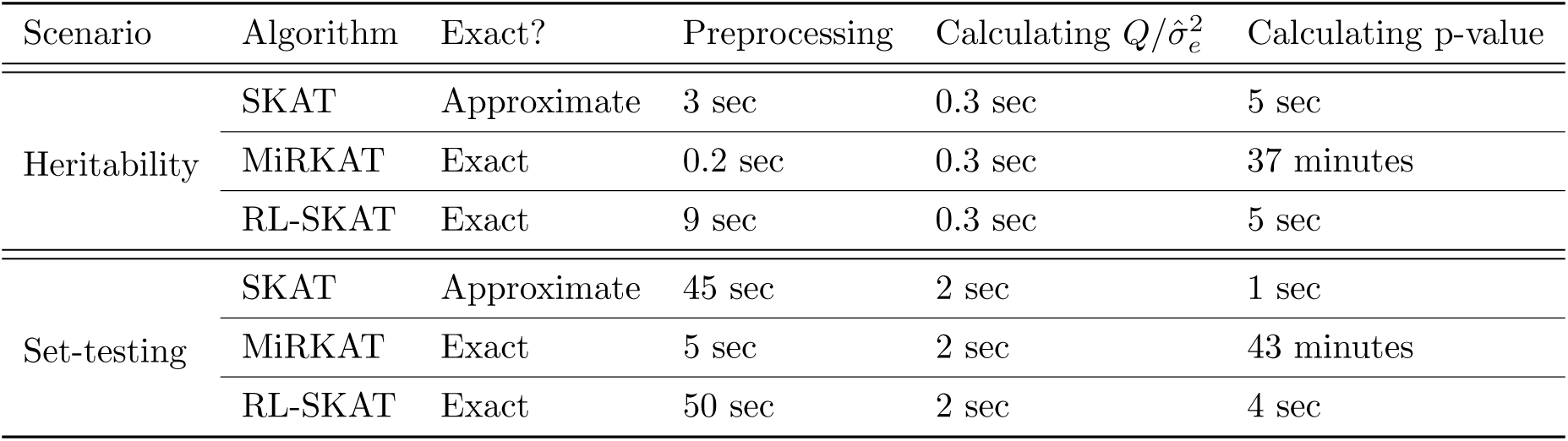
Benchmarks. Benchmark of the performance of different approaches for p-value calculation, applied to the KORA dataset.

## 6 Discussion

In summary, we have shown that the distribution suggested by SKAT to the score test statistic may be very inaccurate. Unlike previous studies, which have noted this discrepancy only in small sample sizes, we have shown that it might occur in large studies as well. We have proposed a computational method to accurately calculate p-values without compromising computational time. Finally, we demonstrated our findings in two datasets.

The exact calculation of p-values can be applied to other variants of the score test; for example, the SKAT-O [16] seeks to find an optimal combination of burden tests and non-burden tests, which amounts to the score test with a certain kernel.

In this work, we focused on the case of a single kernel, and on a continuous phenotype. The extension of this work to multiple kernels (e.g., corresponding to several sets of SNPs) or to binary phenotypes (e.g., case/control studies) is nontrivial, as the null distribution cannot be modeled as a ratio of quadratic forms; see, e.g., [37, 38]. It therefore remains a subject for future work.

## Acknowledgements

R.S. is supported by the Colton Family Foundation. This study was supported in part by a fellowship from the Edmond J. Safra Center for Bioinformatics at Tel Aviv University to R.S. This study makes use of data generated by the Wellcome Trust Case Control Consortium. A full list of the investigators who contributed to the generation of the data is available from www.wtccc.org.uk. Funding for the project was provided by the Wellcome Trust under award 076113. The KORA study was initiated and financed by the Helmholtz Zentrum München German Research Center for Environmental Health, which is funded by the German Federal Ministry of Education and Research (BMBF) and by the State of Bavaria. Furthermore, KORA research was supported within the Munich Center of Health Sciences (MC-Health), Ludwig-Maximilians-Universität, as part of LMUinnovativ.

## Appendix A Proofs

### A.1 Proof of Proposition 1

*Proof.* Note that **S**^2^ = **S** = **S**^⊤^. Under the null, 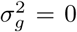 and thus, 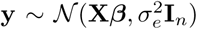. Regressing out the covariates, we get 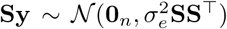 and thus 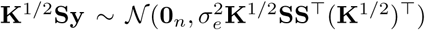. Denote by **U** the matrix whose columns are the eigenvectors of 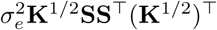, and denote its eigenvalues by 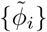. Then, 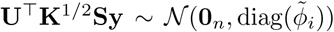. The score test statistic *Q* equals *Q* = ║**K**^1/2^**Sy**║^2^ = ║**U**^⊤^**K**^1/2^**Sy**║^2^, and is therefore distributed

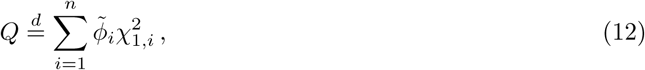

where 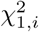 are i.i.d. random variables distributed chi-square with one degree of freedom. It can be seen that 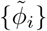 are equivalently the eigenvalues of the matrix 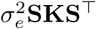, using the fact that for every matrix **A**, **AA**^⊤^ and **A**^⊤^**A** have the same nonzero eigenvalues and that **S** = **S**^⊤^. We divide both the statistic and the eigenvalues by 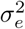; that is, equivalently,

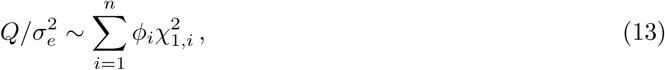

where {*ϕ_i_*} are the eigenvalues of **SKS**^⊤^.

### A.2 Proof of Proposition 2

*Proof.* Assuming 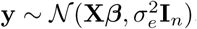, we get 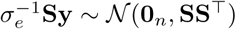. Let 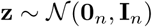; then 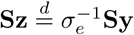. Thus, it follows that

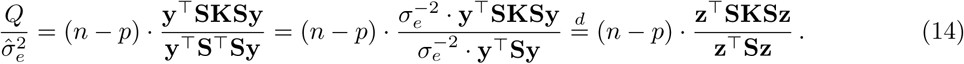

### A.3 Moments of 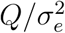 and 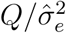

#### Proposition 8.

*Denote the eigenvalues of* **SKS** *by ϕ*_1_*, …, ϕ_n_. Then the expectations of* 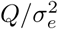 *and* 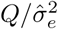 *are identical:*

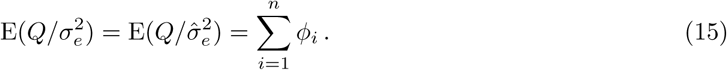

*Their variances are given by:*

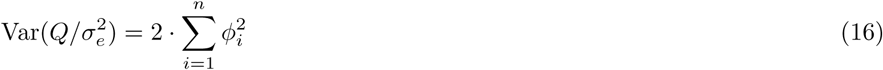

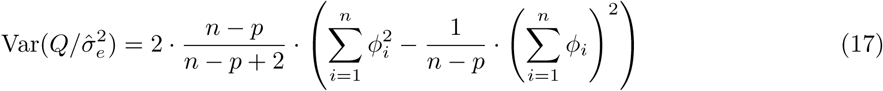

*Proof*. Let 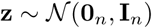, and recall that 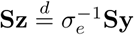. Then, 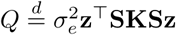. The first two moments of the quadratic form **z**^⊤^**Az**, for a symmetric, positive semi-definite matrix **A**, are readily given [39]. For **A** = **SKS**, they are tr(**SKS**) and 2 · tr(**SKS** · **SKS**), respectively, from which the expressions for 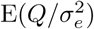 and 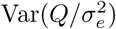 follow.

The first two moments of **z**^⊤^**SKSz**/**z**^⊤^**Sz** are given in [40], and are, respectively,

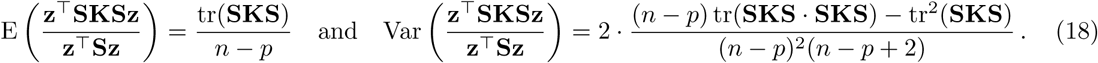

Utilizing Equation (6), the expressions for 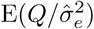 and 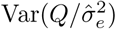 follow.

### A.4 Proof of Proposition 3

*Proof.*

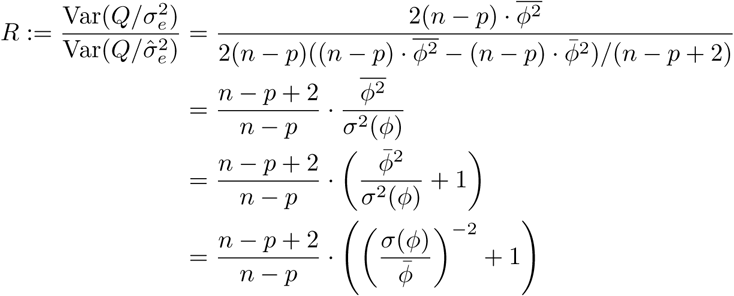

### A.5 Proof of Proposition 4

*Proof.* Let 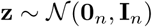, and let **A**, **B** be two symmetric, positive semi-definite matrices. Then (Theorem 3.2d.4 in [39]),

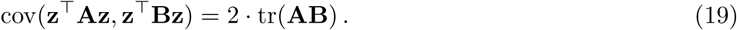

Recall that 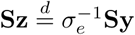. Therefore, ignoring constant factors,

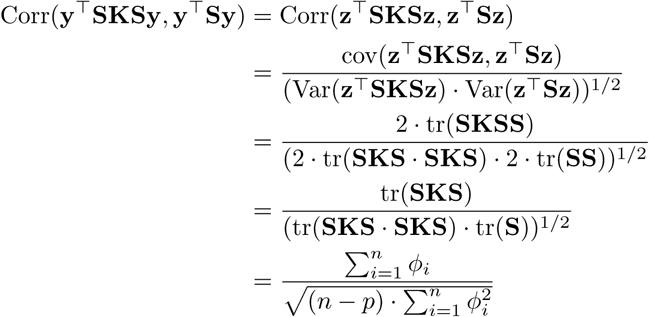

The last equality follows from the fact that **S** is a projection matrix to a *n* − *p* dimensional subspace, and thus its *n* − *p* nonzero eigenvalues are all 1-s, so that tr(**S**) = *n* − *p*. From this,

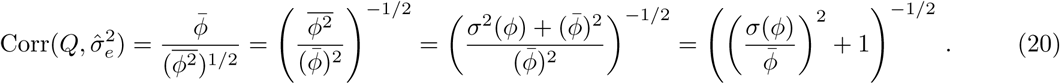

### A.6 Proof of Proposition 5

*Proof.* The condition 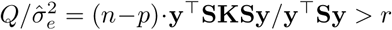 is equivalent to 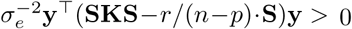. Recall that if 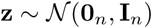, then under the null, 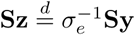. Denote by **A** the matrix whose columns are eigenvectors of **SKS** − *r*/(*n* − *p*) · **S**. Then, also 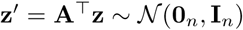. Finally, define 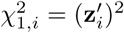. Then,

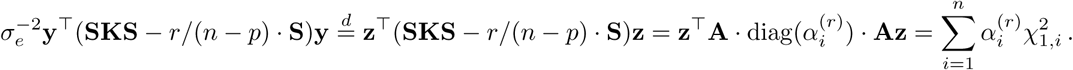

### A.7 Proof of Proposition 6

*Proof.* Denote by ⊕ the direct sum of subspaces. Recall that, from the fundamental theorem of linear algebra, for every symmetric matrix **A**, 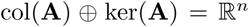. As both **SKS** and **S** are symmetric, we can decompose 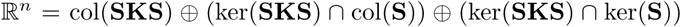. We will construct a basis for each of these subspaces and characterize the operation of **SKS** − *r*/(*n* − *p*) · **S** on them. Note also that from the spectral theorem, there exists an orthonormal basis of eigenvectors of the symmetric matrix **SKS**. In particular, the set of eigenvectors corresponding to nonzero eigenvalues constitutes an orthonormal basis for col(**SKS**).

First, let **v**_1_*, …*, **v**_*k*_ be a basis consisting of such eigenvectors for col(**SKS**), where *k* = rank(**SKS**). Let **v**_*i*_ be an eigenvector of **SKS** corresponding to the eigenvalue *ϕ_i_* > 0. As 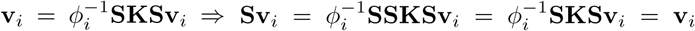, we have that **Sv**_*i*_ = **v**_*i*_, where we used **S**^2^ = **S**. Then, (**SKS** − *r*/(*n* − *p*) · **S**)**v**_*i*_ = (*ϕ_i_* − *r*/(*n* − *p*))**v**_*i*_. Second, let **v**_*k*+1_, …, **v**_*k*+*q*_ be a basis for ker(**SKS**) ∩ col(**S**), where *q* = dim(ker(**SKS**)∩ col(**S**)). For each such **v**_*i*_, (**SKS**−*r*/(*n*−*p*)·**S**)**v**_*i*_ = −*r*/(*n*−*p*)·**v**_*i*_. Finally, let **v**_*k*+*q*+1_,…,**v**_*n*_ be a basis for ker(**SKS**)∩ ker(**S**). Clearly, for each such **v**_*i*_, (**SKS**−*r*/(*n*−*p*)·**S**)**v**_*i*_ = **0**_*n*_.

As before, under the null, 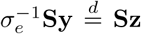, where 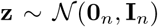. Let **V** be the orthonormal matrix whose columns are **v**_1_*, …*, **v**_*n*_ above; then also 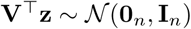. So,

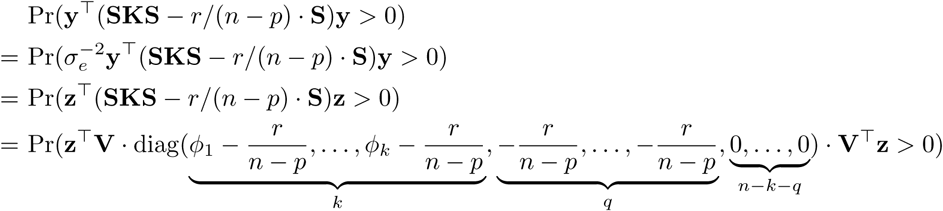

Defining 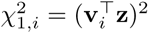, the required result follows.

### A.8 Proof of Proposition 7

*Proof.* Let **B** be the *n*×(*m*+*p*) concatenated matrix **B** = [**SZW**^1/2^, **X**]. Then, rank(**B**) = dim(col(**SKS**)+ col(**X**)) = dim(col(**SKS**) + ker(**S**)), since ker(**S**) = col(**X**). Also note that ker(**SKS**) ∩ col(**S**) = col(**SKS**)^⊥^ ∩ ker(**S**)^⊥^ = (col(**SKS**) + ker(**S**))^⊥^, where ⊥ denotes an orthogonal subspace. Using the above, we can express *q* as:

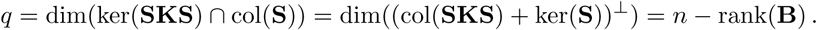

To calculate **SZW**^1/2^, recall that **S** = **I**_*n*_−**X**(**X**^⊤^**X**)^−1^**X**^⊤^, so that **SZW**^1/2^ = **ZW**^1/2^−**X**(**X**^⊤^**X**)^−1^**X**^⊤^**ZW**^1/2^. Calculating **X**^+^ = (**X**^⊤^**X**)^−1^**X**^⊤^ is done once in preprocessing in complexity *O*(*nm*^2^ + *m*^3^). Then, **SZW**^1/2^ can be calculated by multiplying *n* × *p*, *p* × *p*, *n* × *m*, and *m* × *m* matrices, in complexity *O*(*nm*^2^ + *nmp* + *np*^2^) = *O*(*n*(*m* + *p*)^2^). As rank(**SKS**) = rank(**SZW**^1/2^), it can thus be calculated in *O*(*n*(*m* + *p*)^2^). Also, rank(**B**) can be calculated in standard methods (e.g. SVD) in *O*(*n*(*m* + *p*)^2^) as well.

## Appendix B Data preprocessing

### B.1 WTCCC2

UK controls and cases from both UK and non-UK were used. SNPs were removed with > 0.5% missing data, *p* < 0.01 for allele frequency difference between two control groups, *p* < 0.05 for deviation from Hardy-Weinberg equilibrium, *p* < 0.05 for differential missingness between cases and controls, or minor allele frequency < 1%. In all analyses, SNPs within 5M base pairs of the human leukocyte antigen (HLA) region were excluded, resulting in *m*=360,556 SNPs. Finally, from each pair of individuals with relatedness of more than 0.025, one was removed, resulting in *n*=13,950 individuals.

### B.2 KORA

DNA methylation levels were collected using the Infinium HumanMethylation450K BeadChip array (Illumina). Beta Mixture Quantile (BMIQ) [41] normalization was applied to the methylation levels. As described elsewhere [42], genotyping was performed with the Affymetrix 6.0 SNP Array (534,174 SNP markers after quality control), with further imputation using HapMap2 as a reference panel. A total of 657,103 probes remained for the analysis.

